# Exploring Effector Protein Dynamics and Natural Fungicidal Potential in Rice Blast Pathogen *Magnaporthe oryzae*

**DOI:** 10.1101/2024.07.04.602162

**Authors:** Jannatul Ferdausi, Tanjin Barketullah Robin, Sanjida Nasrin, Istiak Ahmed, Tawsif Hossain, Md Mehedi Hasan, Mehrab Hassan Soaeb, Md. Ahsanul Tamim, Nusrat Jahan Yeasmin, Ummay Habiba, Nadim Ahmed, Nurul Amin Rani, Md Shishir Bhuyian, Suvarna N. Vakare, Abu Tayab Moin, Rajesh B. Patil, Mohammad Shahadat Hossain

**Author notes:** These authors contributed equally (Jannatul Ferdausi, Tanjin Barketullah Robin, Sanjida Nasrin, Istiak Ahmed, Tawsif Hossain, Md Mehedi Hasan, Mehrab Hassan Soaeb, Md. Ahsanul Tamim, Nusrat Jahan Yeasmin, Ummay Habiba, Nadim Ahmed, Nurul Amin Rani, Md Shishir Bhuyian, Abu Tayab Moin). **For Correspondence:** Rajesh B. Patil Mohammad Shahadat Hossain Abu Tayab Moin.

## Abstract

Rice blast, caused by *Magnaporthe oryzae*, is a severe agricultural disease leading to significant global economic losses. Genetic and genomic investigations have identified crucial genes and pathways involved in its pathogenesis, particularly highlighting effector proteins like AvrPik variants and MAX proteins. These proteins interact with specific Pik alleles on rice chromosome 11, influencing host immune responses. This study focused on 35 plant-derived metabolites known for their antifungal properties, evaluating their potential as fungicidal agents against M. oryzae. Molecular docking analyses identified Hecogenin and Cucurbitacin E as highly effective binders to MAX40 and APIKL2A proteins, respectively, which are pivotal for fungal virulence and immune evasion. Molecular dynamics simulations further validated strong and stable interactions, affirming the therapeutic potential of these compounds. Additional assessments including Lipinski’s rule of five criteria and toxicity predictions indicated their suitability for agricultural use. These findings underscore the promise of Hecogenin and Cucurbitacin E as lead candidates in developing novel fungicidal strategies against rice blast, offering prospects for enhanced crop protection and agricultural sustainability.

## 1. Introduction

Rice blast disease is a global issue affecting rice-growing regions worldwide (Kirtphaiboon et al., 2021; Shahriar et al., 2020). The fungus, which thrives in warm, humid climates, particularly impacts tropical and subtropical regions where most rice is cultivated. This disease has a substantial financial impact on farmers, costing them billions of dollars annually (Mwangi, 2015; Simkhada & Thapa, 2022). It leads to direct yield lossezs and increased production costs due to the need for fungicides and other control measures. Additionally, rice blast disease reduces rice’s market value, further affecting farmer income (Mew et al., 2004). The production and consumption of rice significantly influence the economies and societies of the Asian subcontinent (Castilla et al., 2021). China, the world’s largest rice producer, contributes over 30% of global rice production, followed by India with around 20% (Bandumula, 2018; Coats, 2003). In many impoverished Asian communities, rice is the primary source of nutritional calories. Beyond providing nutrition, rice farming ensures stability and alleviates poverty. Bangladesh exemplifies the complex relationship between rice cultivation and socioeconomic stability, with rice production contributing about 16% to its GDP and 70% to its agricultural GDP (Dey, 2020).

Global rice production faces numerous challenges, with over 70 infections caused by fungi, bacteria, viruses, or nematodes, 32 of which threaten crop health and yield in Bangladesh alone (Fahad et al., 2019). Rice blast disease, caused by the fungus *Magnaporthe oryzae*, results in significant annual losses in Asia. For example, approximately 564,000 metric tonnes of rice are affected each year in eastern India, especially in upland areas (CO et al., 2011). A severe outbreak occurred during Bangladesh’s Boro season in 2017 and 2018 (Islam et al., 2019).

Managing rice blast disease is challenging due to the high genetic variability of *M. oryzae* (Xu et al., 2019). The fungus can quickly evolve new strains that overcome resistance genes in rice varieties (Jia et al., 2003; Nottéghem, 1993). The disease’s environmental dependence also complicates management, as it is most severe in warm, humid conditions, making it difficult to predict outbreaks and implement preventive measures (Ceresini et al., 2018). Chemical fungicides are often used to control the disease, but natural plant compounds offer a safer and more effective alternative (Lengai et al., 2020).

*M. oryzae* secretes small molecules known as effector proteins during the infection process (Zhang & Xu, 2014). These proteins act like Trojan horses, infiltrating rice plant cells and disrupting their normal functions (Chen et al., 2013; Guo et al., 2019; Qian et al., 2022; Zhang & Xu, 2014). Effector proteins target various cellular processes, suppressing the plant’s defense mechanisms and facilitating fungal colonization. They can inhibit the production of signaling molecules such as salicylic acid and jasmonic acid, which are essential for activating plant defense responses (Liu et al., 2016). Some effectors target enzymes involved in cell wall synthesis and reinforcement, making it easier for the fungus to penetrate and colonize plant tissue (Kubicek et al., 2014). They can also mimic or interfere with plant signaling molecules, tricking the plant’s immune system into suppressing its defenses (Mentlak et al., 2012; Wang et al., 2016).

This study extensively analyzed effector proteins of *M. oryzae*, including AvrPik variants and MAX proteins. The protein structures were refined by removing unwanted ligands, metals, and ions using bioinformatics tools. Thirty secondary metabolites, known for their antifungal properties and derived from plants, were evaluated for their binding affinities with these proteins through molecular docking and dynamics simulations. Fungicide-like properties of these metabolites were assessed according to Lipinski’s rule of five, and general toxicity was evaluated. The results indicated promising binding affinities and acceptable toxicity profiles for the metabolites. However, further in vitro and in vivo studies are necessary to validate and explore these findings.

## 2. Methods

The stepwise methods used in the study are illustrated in **Figure 1**.

**Figure 1:**
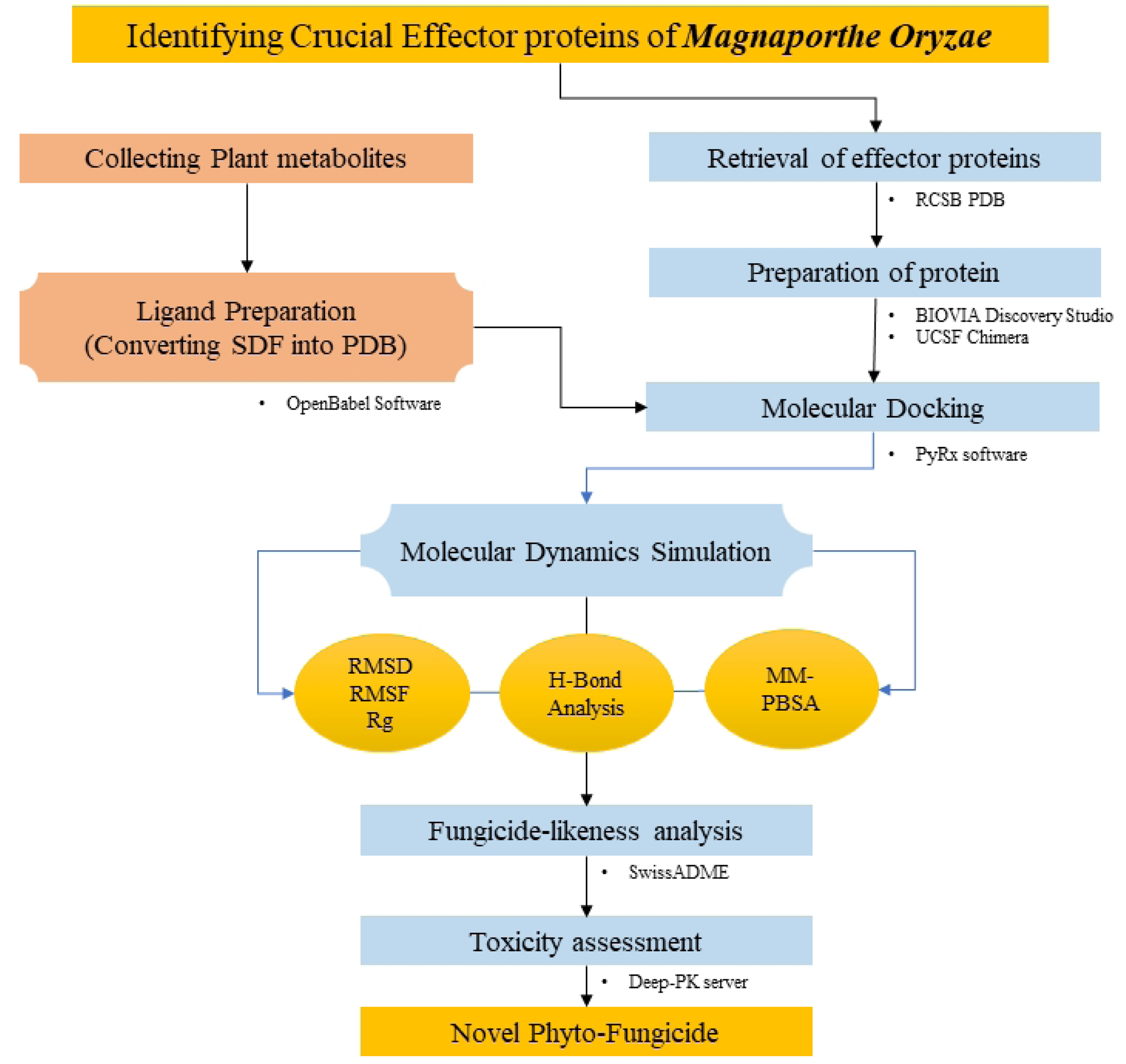
Step-by-step flowchart detailing the methods used in this study

### 2.1. Preparation of the target proteins

Effector proteins of *M. oryzae*, including AvrPik variants and MAX proteins, were selected based on an extensive literature review. The structures of these proteins were obtained from The RCSB Protein Data Bank (Berman et al., 2000; Burley et al., 2021). Subsequently, the target protein structures were refined using the BIOVIA Discovery Studio Visualizer program to remove undesired ligands, metals, and ions (Ghosh et al., 2022). For protein visualization, UCSF Chimera Software (Pettersen et al., 2004) was used.

### Ligand preparation

Thirty secondary metabolites with established antifungal activity from various plant sources were selected through a comprehensive literature review. These substances and inhibitors are potential therapeutic candidates against fungi. Evaluation of their binding affinities utilized reference ligands like strobilurin. The structures of these metabolites, available in SDF (3D) format, were retrieved from the PubChem database (Kim et al., 2019) and converted to PDB format using Open Babel v2.3, a versatile chemical data tool supporting over 110 different formats (O’Boyle et al., 2011), to prepare them for further analysis.

### 2.3. Molecular docking and binding interaction analysis

A flexible docking methodology was employed to investigate the binding affinity between metabolites and proteins (Bappy et al., 2023). The molecular docking of the selected ligands with the target proteins was conducted using AutoDock Vina from the Python Prescription 0.8 (PyRx) package (Suryawanshi et al., 2020). Prior to docking, all compounds and reference drugs underwent minimization and conversion to PDBQT format. The docking procedure itself was executed using Vina Wizard. For the analysis of binding sites and visualization of results, PyMOL v2.0 was utilized to identify polar and non-polar residues and examine the binding locations of specific metabolites (Seeliger & de Groot, 2010; Yuan et al., 2017).

### 2.4. Molecular dynamics simulation analyses

Deeper insights into the binding affinities and possible intricate stabilizing properties of Hecogenine (HEC) and Strobulirin (STR) were obtained from the molecular dynamics (MD) simulations selecting the respective docked complexes of these ligands with rice effector proteins namely ApikL-2a, ApikL-2F, AVR-Pia, AVR-Pib, AVR-Pii, AVR-PikA, AVR-PikC, AVR-PikD, AVR-PikE, AVR-PikF, AVR-PizT, MAX47, MAX60, MAX67. The 100 ns MD simulations were conducted with the Gromacs-2020.4 (Abraham et al., 2015; Berendsen et al., 1995) program on the HPC cluster at Bioinformatics Resources and Applications Facility (BRAF), C-DAC, Pune. The topologies of proteins were built using the parameters from the CHARMM-36 force field (Best et al., 2012; Vanommeslaeghe et al., 2010), while ligand topologies were obtained from the cGenFF server (Vanommeslaeghe et al., 2010). The respective complexes were held in a dodecahedron unit cell such that the edges of the system were 1 nm away from the edges of the box. Subsequently, the systems were solvated with water molecules employing the TIP3P water model (Jorgensen & Madura, 1983) and neutralized by adding sodium and chloride counter-ions to achieve 0.15 mole concentration. Energy minimization was performed to relieve the steric strains with the steepest descent algorithm until the force constant reached the 100 kJ mol-1 nm-1 threshold. The systems were then equilibrated at constant temperature and pressure conditions, NVT and NPT conditions, at 300 K temperature and 1 atm pressure condition using a modified Berendsen thermostat (Bussi et al., 2007) and barostat (Berendsen et al., 1984), respectively, for 1 ns each. The 100 ns MD simulations were performed on the equilibrated systems where the temperature conditions of 300 K were achieved with a modified Berendensen thermostat, and pressure conditions of 1 atm were achieved with the Parrinello-Rahman barostat (Parrinello & Rahman, 1981). The covalent bonds in the systems were restrained with the LINCS algorithm (Hess et al., 1997). The long-range electrostatic energies were computed with the Particle Mesh Ewald (PME) method (Petersen, 1995) at a cut-off of 1.2 nm. After removing the periodic boundary conditions, the trajectories of each complex were analyzed for the root mean square deviations (RMSD) in the backbone atoms of proteins. The RMSD in ligand atoms was analyzed by selecting protein backbone atoms for the most minor square fitting and ligand atoms for RMSD calculations, effectively showing the relative positions of ligands compared to initial position during the simulation.

Further, the root mean square fluctuation (RMSF) in the side chain atoms was analyzed. The radius of gyration (Rg) of the protein structure from its center of mass was analyzed to investigate the stability and compactness of the corresponding systems. During the simulation, the hydrogen bonds between the ligand and the protein residues were analyzed and visually inspected in the trajectories extracted at steps 0, 25, 50, 75, and 100 ns. Molecular Mechanics General Born surface area and surface area solvation (MM/GBSA) (Valdés-Tresanco et al., 2021) calculations were performed on the trajectories isolated from the simulation period 75 to 100 ns at each 100 ps time step. The entropic energies were accounted for to obtain each complex’s binding free energies (ΔGbinding kcal/mol). The protein-ligand structures were rendered in PyMOL (Sufyan et al., 2021) and graphs were obtained from XMGRACE (Turner, 2005)

### 2.5. Fungicide-like properties and toxicity analysis

The fungicide-likeness of natural compounds was assessed using Lipinski’s rule of 5, a well-established criterion for drug-likeness, given the absence of similar criteria for fungicides (Walters, 2012). The SwissADME server (http://www.swissadme.ch/) was utilized to evaluate the fungicidal qualities of the top metabolites (Daina et al., 2014, 2017), providing predictions based on the compounds’ characteristics related to Lipinski’s Rule (Daina & Zoete, 2016; Khan et al., 2022). Afterwards, the general toxicity of the identified compounds was forecasted using the pkCSM web-based server (https://biosig.lab.uq.edu.au/pkcsm/) (Pires et al., 2015).

## 3. Result

### 3.1. Preparation of the target proteins

Through an extensive literature review, fourteen crucial effector proteins necessary to inhibit M. oryzae, the causative agent of rice blast disease, were identified. These proteins, including APIKL2A, APIKL2F, AVRPIA, AVRPIB, AVRPII, AVRPIKA, AVRPIKC, AVRPIKD, AVRPIKE, AVRPIKF, AVRPIZT, MAX60, MAX47, and MAX67, were sourced from the Protein Data Bank with the following PDB IDs: 7NLJ, 6FUD, 7QPX, 7BNT, 6G11, 7B1I, 6Q76, 5Z1V, 7PP2, 2LW6, 7NMM, 72JY, 7ZK0, 7ZKD respectively. Afterwards, The obtained proteins were processed using BIOVIA Discovery Studio to remove undesired macromolecule ligands, water molecules, and heteroatoms from their structures. Subsequently, molecular graphics and analyses were conducted to explore their properties and potential for inhibiting M. oryzae. The proteins were visualized using PyMOL software, as depicted in **Figure 2**.

**Figure 2:**
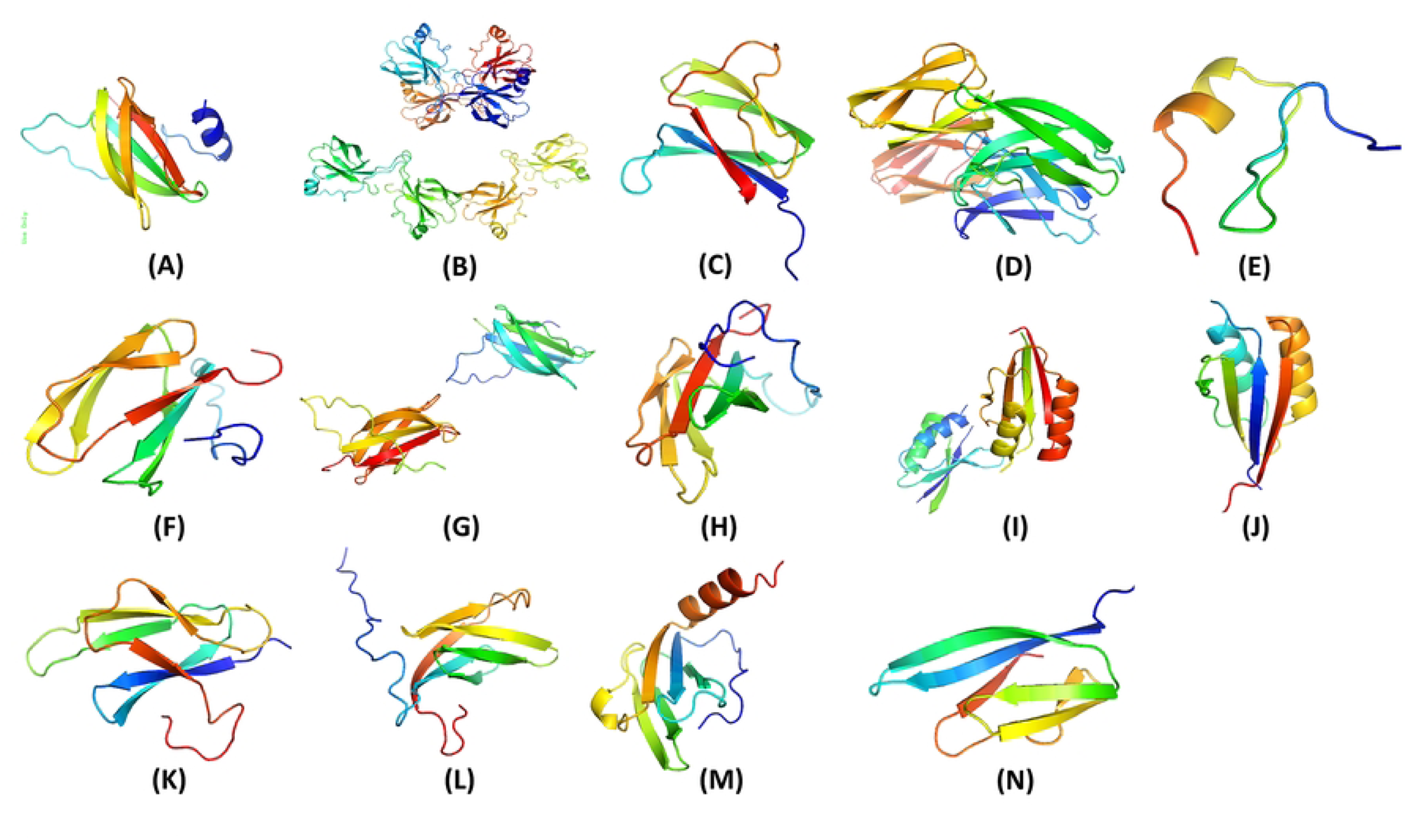
Tertiary structure of A)APIKL2A, B)APIKL2F, C)AVRPIA, D)AVRPIB, E)AVRPII, F)AVRPIKA, G)AVRPIKC, H)AVRPIKD, I)AVRPIKE, J)AVRPIKF, K)AVRPIZT, L)MAX60, M)MAX47, N)MAX67

### 3.2. Ligand Preparation

A thorough literature review was conducted to compile a list of 35 metabolites (**Table** S1) derived from diverse plant species, recognized for their antifungal properties. These metabolites, including inhibitors and chemicals, hold promise as therapeutic agents for combating various fungal infections. 3D structures of these 30 metabolites were obtained from the PubChem database in SDF format, and subsequently converted to PDB format using Open Babel v2.3 software. The same method was applied to convert the reference compound strobilurin into PDB format.

### 3.3. Molecular Docking Studies and Binding site analysis

Molecular docking experiments were performed using 14 effector proteins as receptors and 35 plant metabolites (**Table** S2) as ligands. Notably, Hecogenin and Cucurbitacin E exhibited the highest and most consistent binding affinities across all proteins. Hecogenin achieved the best docking energy of -8.8 kj/mol with protein APIKL2A, binding to residues LYS29, TYR25, ILE108, LYS109, and GLY92. Meanwhile, Cucurbitacin E demonstrated a maximum docking energy of -8.0 kj/mol with MAX40 protein, interacting with residues ARG-65, TRP-54, CYS-108, and GLY-106. Despite showing varied energy levels with some proteins, Cucurbitacin E consistently displayed superior binding energy compared to other metabolites. Several metabolites were excluded due to altered drug effects and toxicity. **Table** 1 illustrates that Hecogenin and Cucurbitacin E outperformed the reference metabolite Strobilurin. The binding sites, crucial for complex formation, are depicted in **Figure**s 3 - 5.

**Table 1:**
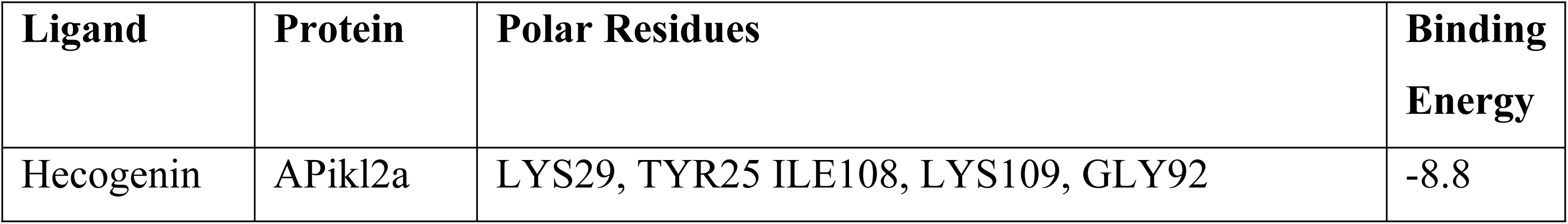
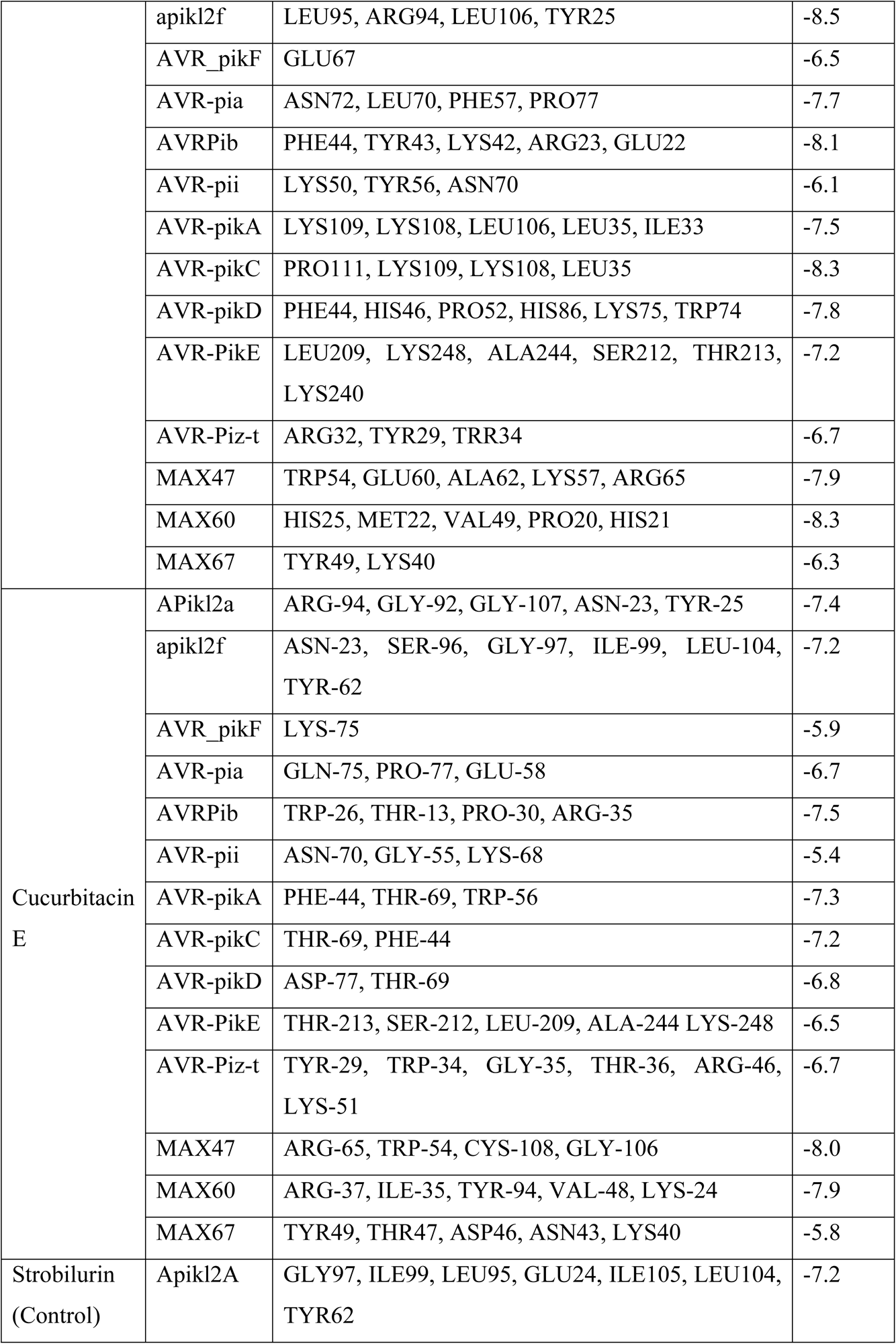
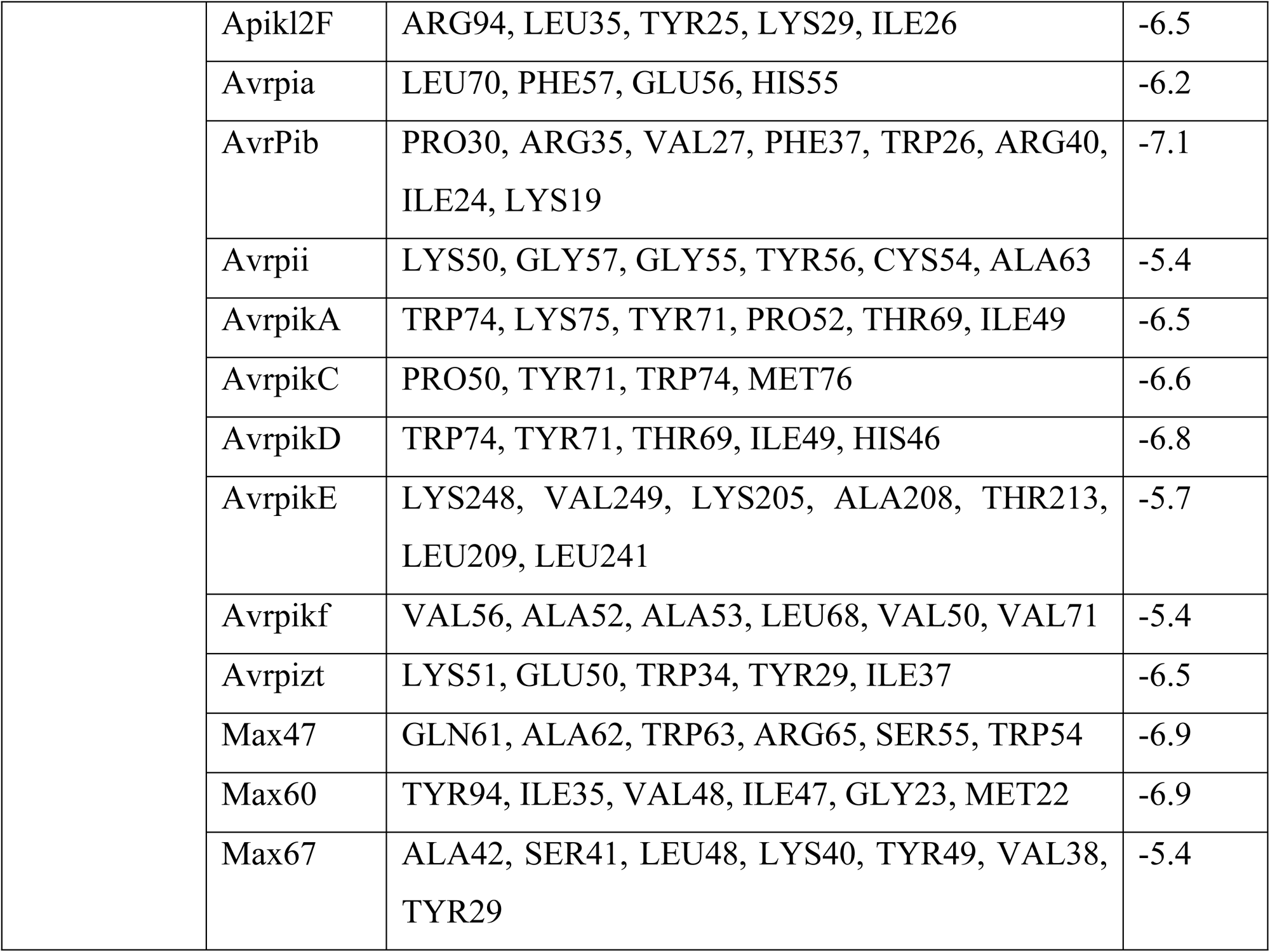
Binding energy and binding site analysis of best complexes.

**Figure 3.**
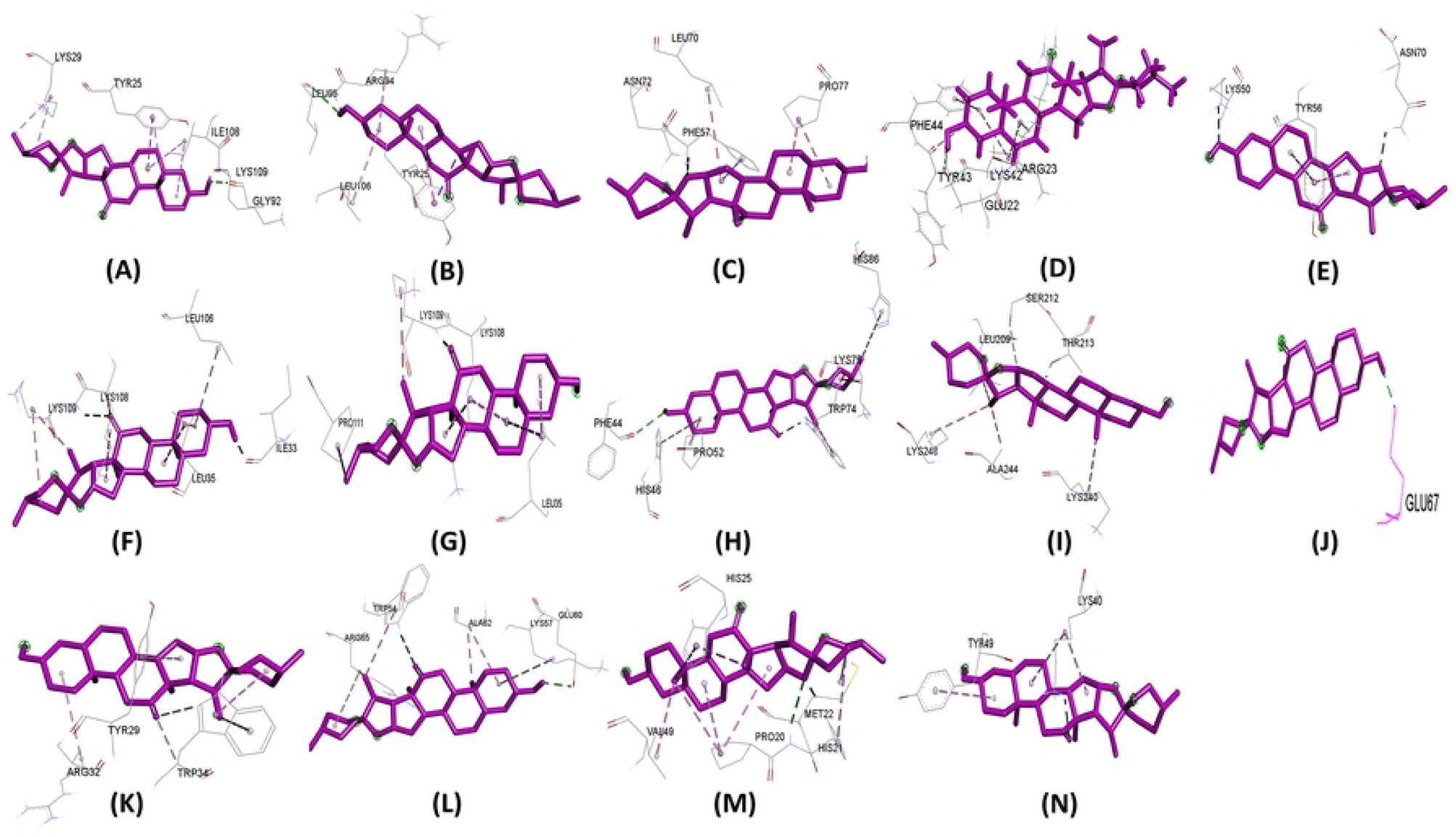
Binding site of ligand Hecogenin with protein A)APIKL2A, B)APIKL2F, C)AVRPIA, D)AVRPIB, E)AVRPII, F)AVRPIKA, G)AVRPIKC, H)AVRPIKD, I)AVRPIKE, J)AVRPIKF, K)AVRPIZT, L)MAX60, M)MAX47, N)MAX67

**Figure 4:**
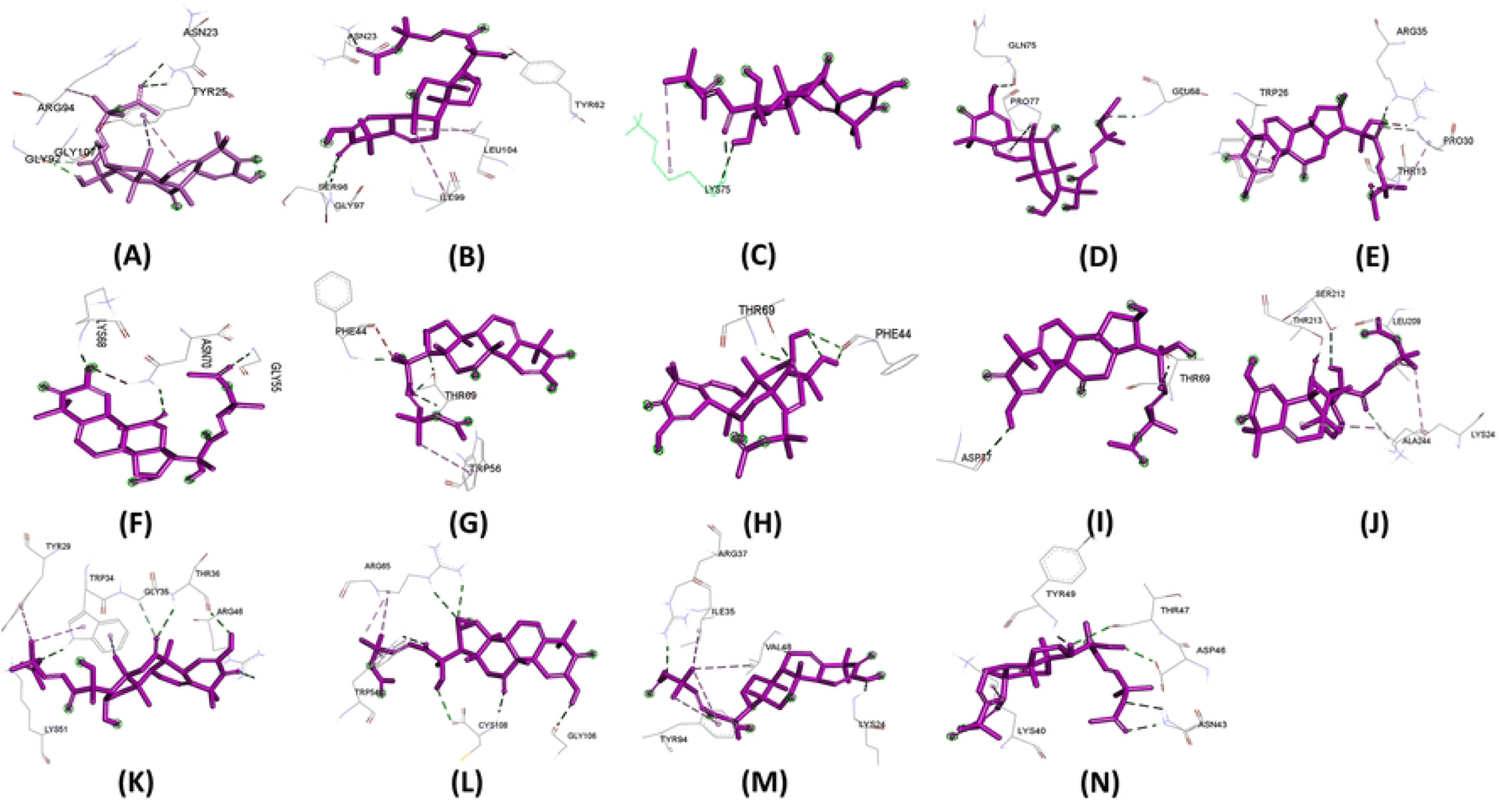
Binding site of ligand Cucurbitacin E with protein A)APIKL2A, B)APIKL2F, C)AVRPIA, D)AVRPIB, E)AVRPII, F)AVRPIKA, G)AVRPIKC, H)AVRPIKD, I)AVRPIKE, J)AVRPIKF, K)AVRPIZT, L)MAX60, M)MAX47, N)MAX67

**Figure 5:**
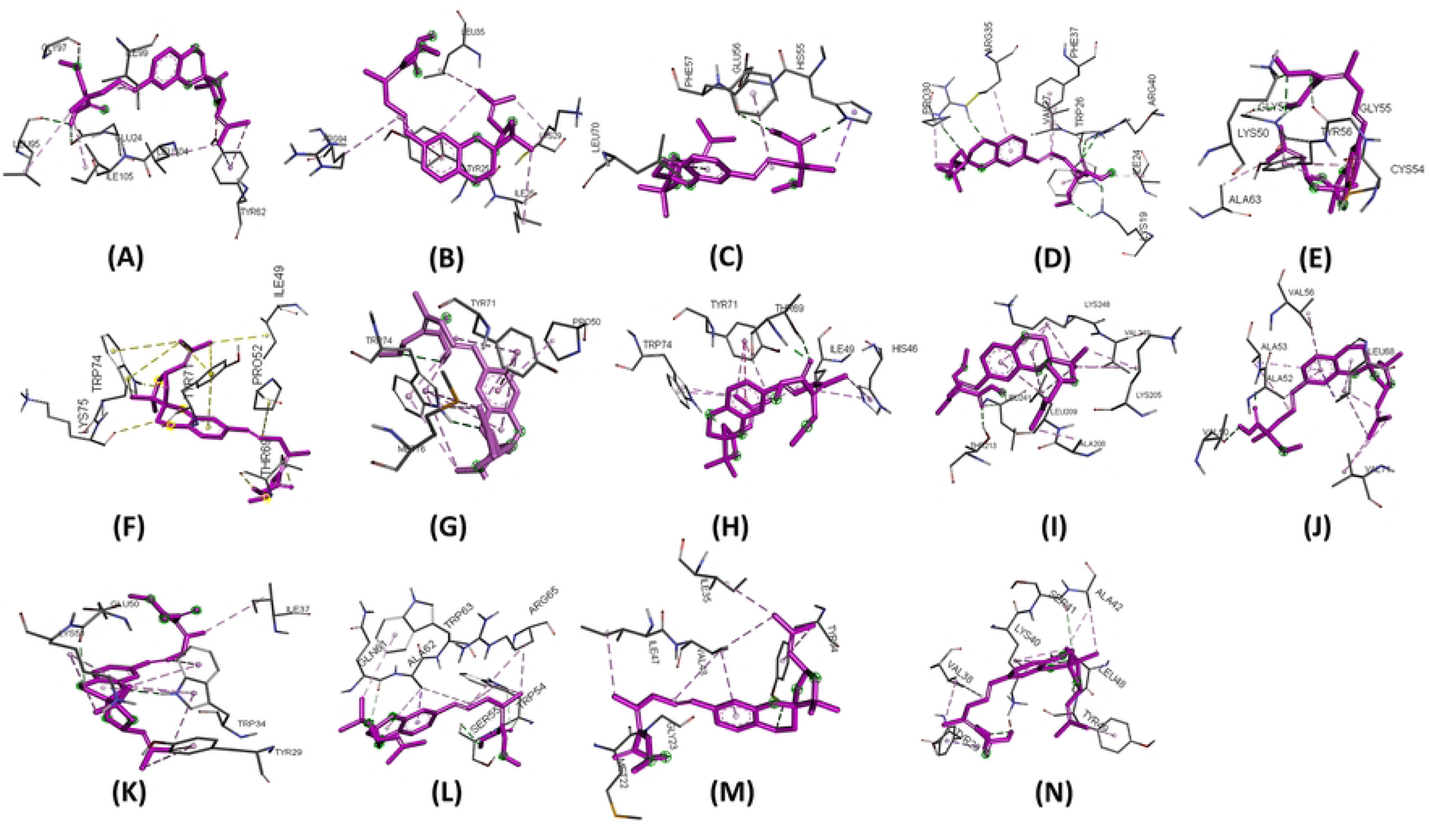
Binding site of ligand Strobilurin (control) with protein A)APIKL2A, B)APIKL2F, C)AVRPIA, D)AVRPIB, E)AVRPII, F)AVRPIKA, G)AVRPIKC, H)AVRPIKD, I)AVRPIKE, J)AVRPIKF, K)AVRPIZT, L)MAX60, M)MAX47, N)MAX67

### 3.4. Molecular dynamics simulation studies

#### 3.4.1. Root mean square deviation

The results of RMSD in the backbone atoms of rice effector proteins showed that the proteins ApikL-2a, ApikL-2f, AVR-Pib, AVR-PikA, AVR-PikC, and MAX67 stabilized reasonably, and the RMSD converged stably throughout the simulation with RMSD below 0.2 nm (**Figure** 6). However, ApikL-2f in complex with STR showed significant deviations after around 80 ns, reaching RMSD of a maximum of 0.3 nm. In the case of AVR-Pia, the complex with HEC showed reasonably stable RMSD compared to the complex with STR, which showed significantly larger RMSD after around 45 ns until the end of the simulation. The RMSD in AVR-Pii was significantly larger for both the complexes with HEC and STR. The AVR-PikC complex with HEC showed significant deviations after around 40 ns until the end of the simulation compared to relatively stable and converged RMSD in the complex with STR. Both the complexes of AVR-PikD showed reasonably stable RMSD with an average of around 0.3 nm. The complexes of AVR-PikE and AVR-PikF with HEC and STR showed substantial deviations throughout the simulation period. However, the AVR-PikE complex with HEC showed slightly lower RMSD stabilizing after around 40 ns to an average of around 0.3 nm.

**Figure 6.**
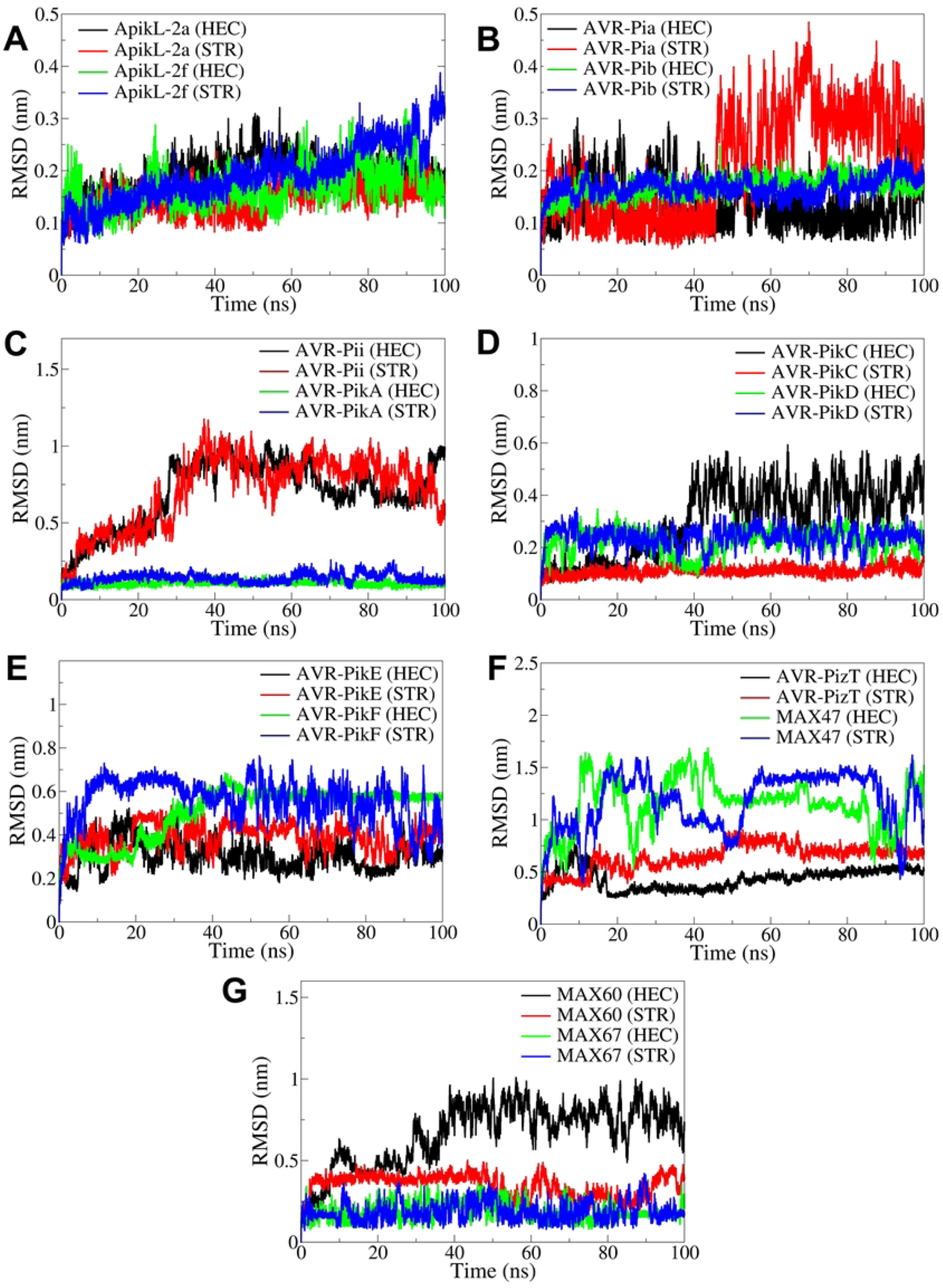
The RMSD analysis in protein backbone atoms. RMSD in backbone atoms of A) ApikL-2a and ApikL-2f, B) AVR-Pia and AVR-Pib, C) AVR-Pii and AVR-PikA, D) AVR-PikC and AVR-PikD, E) AVR-PikE and AVR-PikF, F) AVR-PizT and MAX47, and G) MAX60 and MAX67.

AVR-PizT in complexes with HEC showed comparably lower RMSD than in complexes with STR. The complexes of MAX47 showed substantially more significant deviations than those with HEC and STR. In the case of MAX60, the RMSD in the complex with STR was slightly lower, with an average of around 0.4 nm, compared to the complex with HEC with substantially larger RMSD, reaching a maximum of around 0.9 nm.

The results of RMSD in ligand atoms showed that the ligands in complex with ApikL-2a, AVR-Pib, AVR-PikA, AVR-PizR, and MAX67 had significant deviations relative to the backbone atoms (**Figure** 7). In the case of ApikL-2f, the average RMSD for HEC and STR was around 1 nm. The RMSD was slightly higher and was around 2 nm for the ligands in complex with AVR-Pia and AVR-Pii. In the case of ACR-PikC compared to STR, the ligand HEC had significantly higher RMSD, where RMSD for STR was around 1 nm. In the case of AVR-PikE, both the ligands had lower and almost stable RMSD with an average of around 0.5 nm. The RMSD was relatively stable until around 80 ns in STR atoms in complex with MAX47, while in HEC, the RMSD stabilized to an average of around 0.2 nm after around 40 ns. The RMSD in STR atoms in the complex with MAX60 was relatively stable and lowest, with an average of around 0.5 nm, while the RMSD in HEC was significantly higher, with an average of around 2 nm.

**Figure 7.**
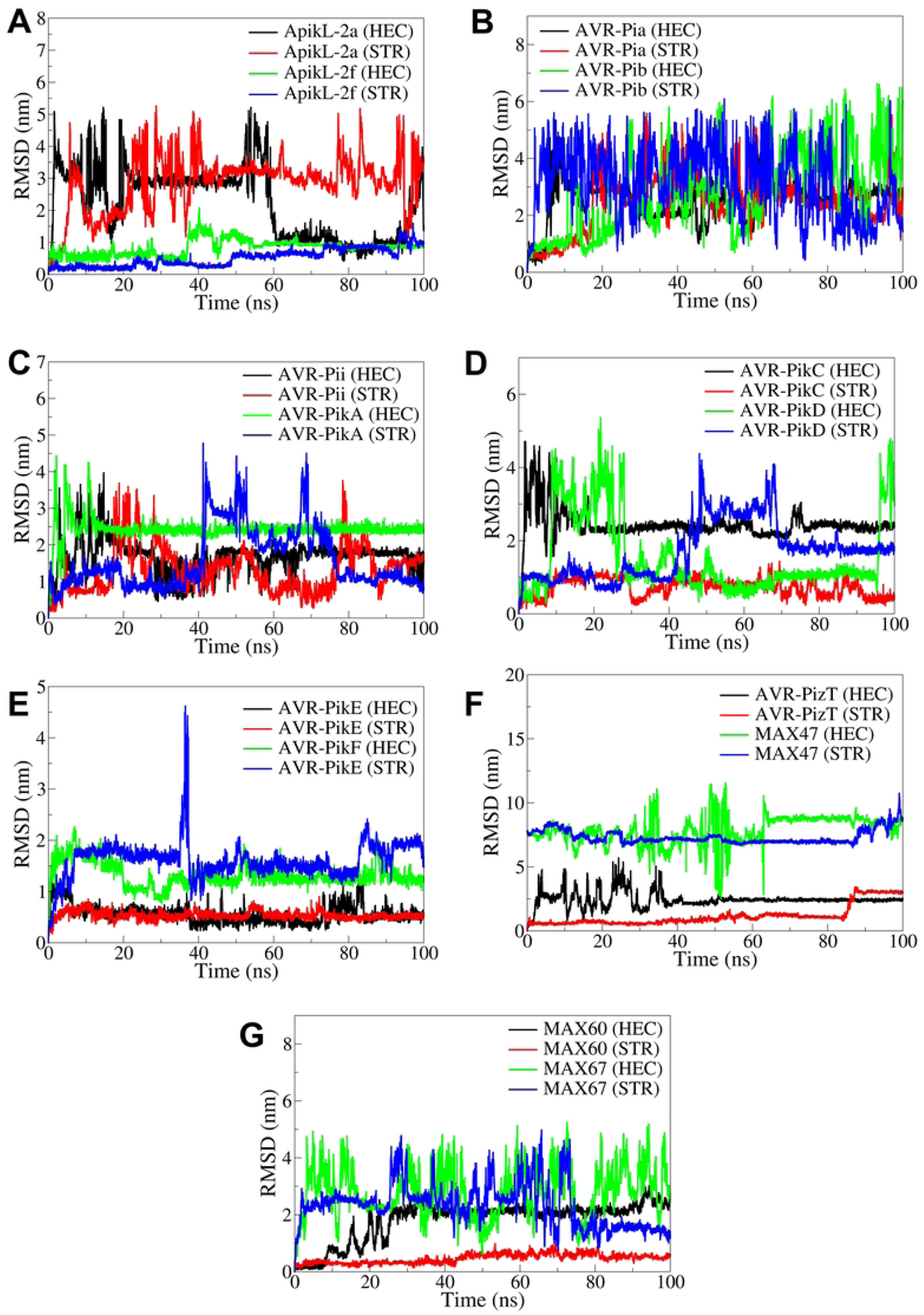
The RMSD analysis in UNK atoms. RMSD in UNK atoms in complexes of A) ApikL-2a and ApikL-2f, B) AVR-Pia and AVR-Pib, C) AVR-Pii and AVR-PikA, D) AVR-PikC and AVR-PikD, E) AVR-PikE and AVR-PikF, F) AVR-PizT and MAX47, and G) MAX60 and MAX67.

#### 3.4.2. Root mean square fluctuation

The RMSF analysis results suggested that for the RMSF in side chain atoms of respective proteins in the complex of both the ligands with ApikL-2a, ApikL-2f, AVR-PikA, AVR-PikC, AVR-PikD, AVR-PikE, AVR-PikF, AVR-PizT, MAX-60, and MAX67 were below 0.4 nm, excluding the higher RMSD in few of the terminal residues (**Figure** 8). The RMSF in the complexes with AVR-Pii and MAX47 was significantly higher. In the case of complexes with ApikL-2a and ApikL-2f, the residues in the range 25-50 showed significant RMSF, whereas the remaining residues showed stability below 0.2 nm RMSF. The complex of AVR-Pib with STR showed significantly higher RMSD in the residues in the range 40-50. The complexes with AVR-PikA, AVR-PikC, and AVR-PikD showed significant fluctuations in the residues in the range 25-50, while the fluctuation in the other residues, except terminal residues, was well below 0.2 nm. The RMSF was found considerably stable in AVR-PizT, MAX-60, and MAX67 complexes.

**Figure 8.**
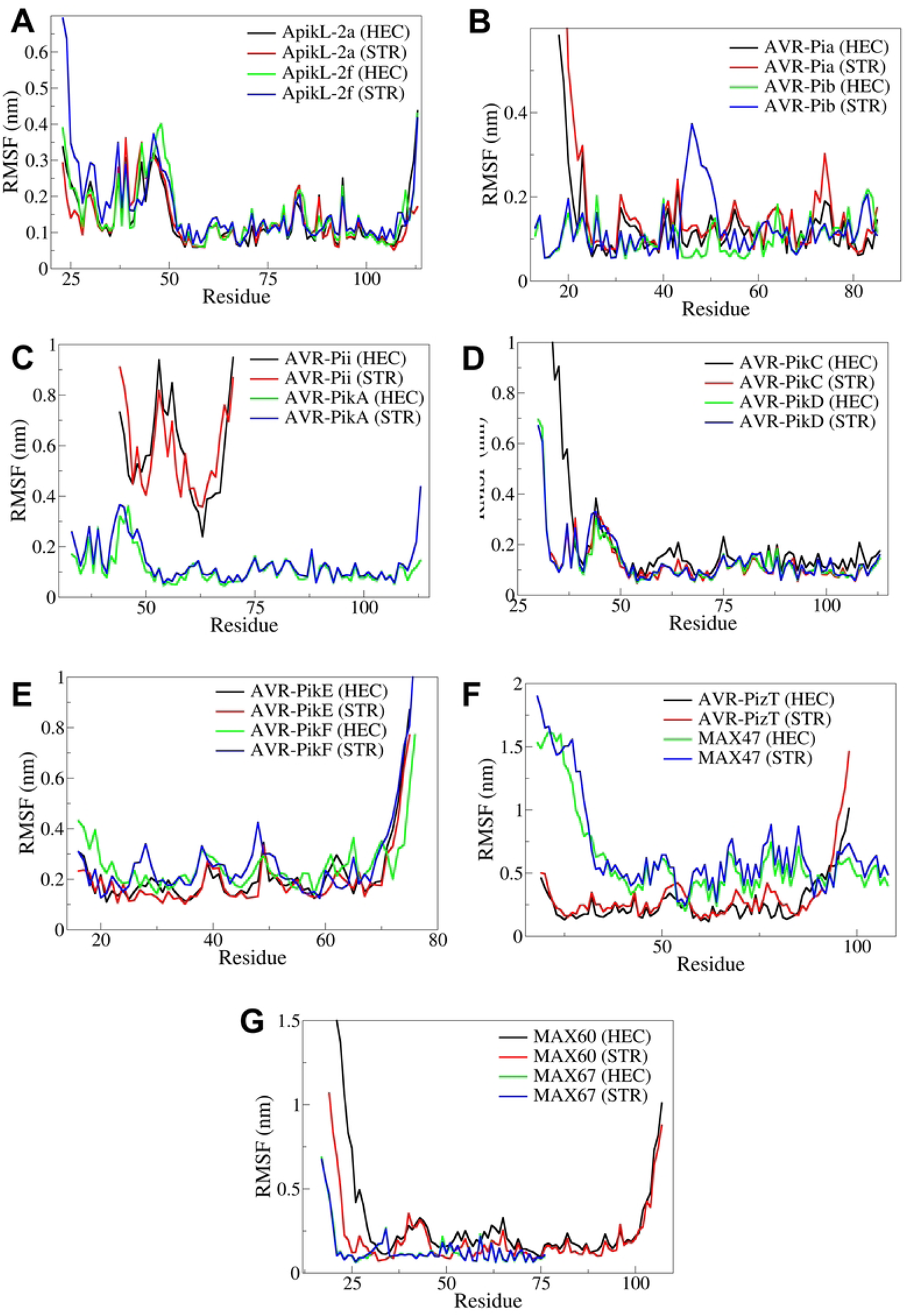
The RMSF analysis in the complexes of A) ApikL-2a and ApikL-2f, B) AVR-Pia and AVR-Pib, C) AVR-Pii and AVR-PikA, D) AVR-PikC and AVR-PikD, E) AVR-PikE and AVR-PikF, F) AVR-PizT and MAX47, and G) MAX60 and MAX67.

#### 3.4.3. Radius of gyration

The radius of gyration results showed varied Rg in different complexes. The Rg in ApikL-2a and ApikL-2f were 1.3 to 1.4 nm (**Figure** 9). However, the ApikL-2f showed slight deviations after around 50 ns in both the complexes with HEC and STR. The Rg in AVR-Pia was almost stable for both the complexes with HEC and STR, while the Rg in AVR-Pib was lower and stable. The complexes of AVR-Pii showed significant deviations in Rg for both the complexes after around 20 ns simulation period. Meanwhile, the complexes of AVR-PikA showed stable Rg. In the case of AVR-PikC, the complex with HEC showed significantly higher Rg after around 40 ns, compared to the stable Rg in the complex with STR. Meanwhile, the Rg in both AVR-PikD complexes was stable. The Rg in AVR-PikE complexes was lower and stable throughout the simulation. Meanwhile, the Rg in AVR-PikF significantly deviated throughout the simulation for the complex with STR compared to the stable Rg in the complex with HEC. The Rg in AVR-PizT was lower and stable in both complexes, whereas the Rg in both complexes of MAX47 deviated significantly throughout the simulation. The Rg in the MAX60 complex with STR was stable compared to the complex with HEC, which showed significant deviations after around 40 ns simulation period. The Rg in both the complexes of MAX67 were lower and stable in both the complexes.

**Figure 9.**
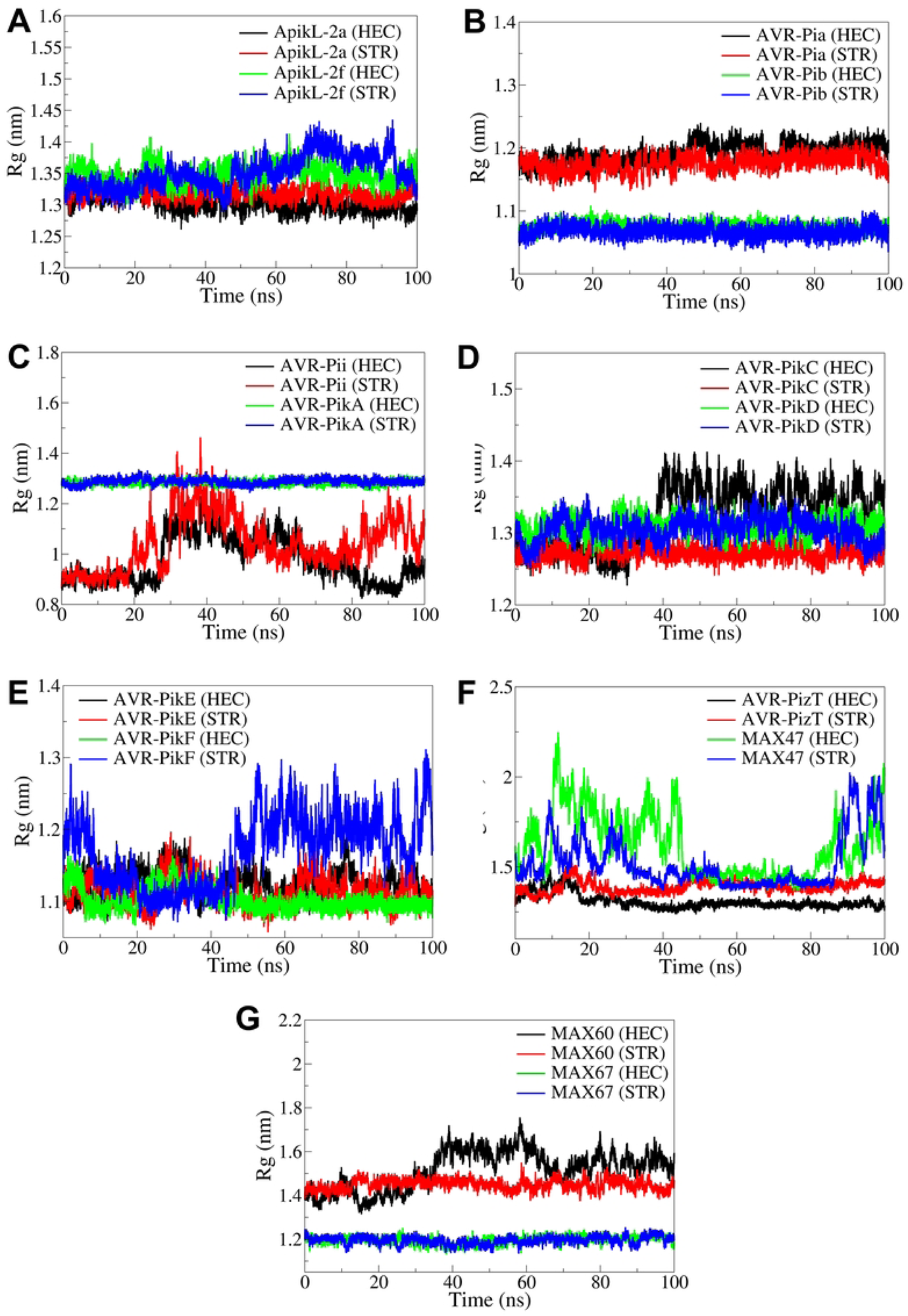
The Rg analysis in the complexes of A) ApikL-2a and ApikL-2f, B) AVR-Pia and AVR-Pib, C) AVR-Pii and AVR-PikA, D) AVR-PikC and AVR-PikD, E) AVR-PikE and AVR-PikF, F) AVR-PizT and MAX47, and G) MAX60 and MAX67.

#### 3.4.4. Hydrogen bond analysis

All the complexes investigated showed at least one hydrogen bond consistently formed throughout the simulation period (**Figure** 10). A maximum of two frequent hydrogen bonds were observed in the complexes of ApikL-2a and APikL-2f, where STR showed more hydrogen bonds for a brief period of 25 to 35 ns in the case of APikL-2f. Compared to the complex of AVR-Pia with STR, the complex with HEC showed more number and more frequent hydrogen bonds. Similarly, in the AVR-Pib complex with STR, HEC showed more and more frequent hydrogen bonds. In the case of AVR-Pii complexes, HEC showed more hydrogen bonds than STR, while in the case of complexes with AVR-PikA, STR showed more frequent hydrogen bonds. In the case of complexes of AVR-PikC and AVR-PikD, both the ligands formed a maximum of two hydrogen bonds frequently, except the STR in AVR-PikD, which showed a number of hydrogen bonds during the first 40 ns simulation period. Both the ligands formed two hydrogen bonds frequently in the complexes of AVR-PikE and AVR-PikF. In the case of AVR-PizT, both the ligands formed a maximum of two hydrogen bonds quite frequently throughout the simulation. The ligand STR formed more consistent and frequent hydrogen bonds with MAX47 than HEC. The ligands formed a maximum of three hydrogen bonds during the first 25 ns simulation period in MAX60 and MAX67 complexes. However, after 25 ns, the ligand STR showed consistent hydrogen bonds compared to HEC in these complexes.

**Figure 10.**
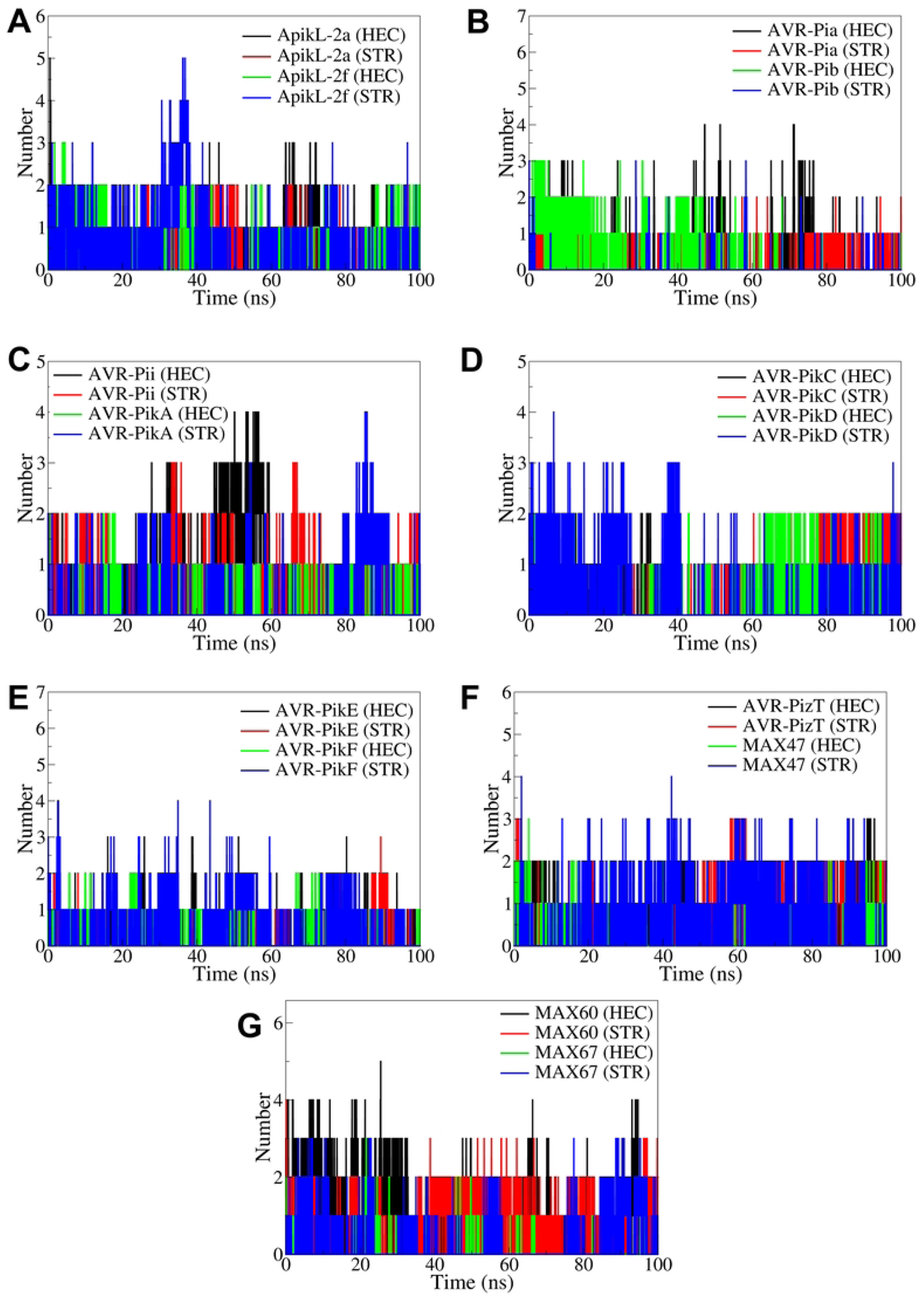
Hydrogen bond analysis in the complexes of A) ApikL-2a and ApikL-2f, B) AVR-Pia and AVR-Pib, C) AVR-Pii and AVR-PikA, D) AVR-PikC and AVR-PikD, E) AVR-PikE and AVR-PikF, F) AVR-PizT and MAX47, and G) MAX60 and MAX67.

In the ApikL-2a complex, the ligand HEC formed hydrogen bonds with Lys109 and Gly92 at the simulation time intervals 0 and 25 ns (**Figure** S1). However, the 50, 75, and 100 trajectories showed no hydrogen bonds. The ligand STR showed the hydrogen bonds with Gly97 and Leu95 were shown only in 0 and 25 ns trajectories. In the case of the ApikL-2f complex, the ligand HEC formed hydrogen bonds with Leu95 in the 0 and 25 ns trajectories, while with Met108 in the 75 ns trajectory, and Arg94 and Leu35 in the 100 ns trajectory. The complex with STR showed a hydrogen bond with Asn23 in 0 and 25 ns trajectories, while the 75 ns trajectory showed a hydrogen bond with Glu24. AVR-Pia only equilibrated from showed a hydrogen bond between HEC and Asn72 (**Figure** S2). At the same time, STR formed hydrogen bonds with Glu58, Glu56, and His55 in an equilibrated trajectory, with residue Thr47 at 25 ns and residue Agr43 at 75 ns. In the case of AVR-Pib, only the equilibrated trajectory showed hydrogen bonds between HEC and Lys60, while the STR showed hydrogen bonds with Lys19 in equilibrated and 25 ns trajectory. In the complex of AVR-Pii, the ligand HEC showed no hydrogen bonds with Asp59, Cys51, and Cys69 in the 50 ns trajectory, while the trajectories at other time steps showed no hydrogen bonds (**Figure** S3). The ligand STR showed hydrogen bonds with Gly57, Thr56, and Lys68 in the equilibrated trajectory and Trp74 and Thr69 in the 25 ns trajectory. The 50 and 75 ns trajectories showed hydrogen bonds between STR and Ser52 and Asn70, respectively. The 100 ns trajectory showed hydrogen bonds with Tyr64 and Asp62. In the case of the complex with AVR-PikA, the ligand HEC showed no hydrogen bonds in the extracted trajectories. The ligand STR showed hydrogen bonds with Trp74 and Thr69 at 0 ns and 25 ns and residues Asn46 and Asp45 at 100 ns.

In the AVR-PikC complex, the ligand HEC formed hydrogen bonds with Lys109 in equilibrated and 25 ns trajectory and Thr69 in 100 ns trajectory (**Figure** S4). Meanwhile, the ligand STR only showed a hydrogen bond with Asn46 in the 75 ns trajectory. In the case of the AVR-PikD complex, the ligand HEC formed hydrogen bonds with Lys75 and Phe44 in the equilibrated and 25 ns trajectory and with Thr69 in the 75 ns trajectory. The ligand STR showed hydrogen bonds with Trp74, Thr69, and Asn42 residues in equilibrated and 25 ns trajectories and Asn83 in 50 ns trajectory. In the case of AVR-PikE, only the ligand HEC showed a hydrogen bond with Gly235 in an equilibrated and 25 ns trajectory (**Figure** S5). In the case of the complex of AVR-PikF, the ligand HEC formed a hydrogen bond with Ser18 in 25 and 50 ns trajectories. Meanwhile, the ligand STR formed hydrogen bonds with Val50, Ala52, and Leu68 in equilibrated and 25 ns trajectories and with Asn51 residue in 75 ns trajectory.

In the case of AVR-PizT complexes, the ligand HEC formed a hydrogen bond with residue His33 in equilibrated and 25 ns trajectories and with Leu58 in 75 and 100 ns trajectories (**Figure** S6). Meanwhile, the ligand STR formed a hydrogen bond with Arg46 in equilibrated and 25 ns trajectories and Val96 and Val21 residues in 100 ns trajectory. The equilibrated and 25 ns trajectories of the MAX47 complex with HEC showed hydrogen bonds with Trp63 and Trp54. The ligand STR showed a hydrogen bond with Ile53 in equilibrated and 25 ns trajectories, Trp63 in 50 ns trajectory, and Lys57 in 75 ns trajectory. In the case of complexes of MAX60, the ligand HEC formed a hydrogen bond with Asp56 in equilibrated and 25 ns trajectories, with Asn101 in 50 ns trajectory, and with Asn101 and Gln97 residues in 100 ns trajectory (**Figure** S7). The ligand STR formed hydrogen bonds with Tyr94 and Gly23 in equilibrated and 25 ns trajectories, with Gln97 in 50 ns trajectory and Gln45 in 100 ns trajectory. In the complex of MAX67, the ligand HEC formed a hydrogen bond with Thr47 in equilibrated and 25 ns trajectory, while STR formed a hydrogen bond with Lys40 in equilibrated and 25 ns trajectory.

#### 3.4.5. MM-GBSA calculations

The results of MM-GBSA calculations are given in **Table 2**. The ligand HEC showed comparably more favorable binding affinities compared to STR in terms of lower ΔG_binding_ in the complexes of ApikL-2a, AVR-Pia, AVR-Pii, AVR-PikA, AVR-PikC, AVR-PikD, AVR-PikF, AVR-PizT, and MAX47. In all these complexes, either due to favorable entropic energies, due to lower van der Waals energies, or due to higher electrostatic energies, the relative binding energies (ΔTOTAL) and the binding free energy (ΔG_binding_) were lower for HEC, notably, in the case of ApikL-2a, AVR-Pia, and MAX47, the ligand HEC had comparably and significantly lower ΔG_binding_. However, in the complexes with AVR-Pib, AVR-PiE, MAX60, and MAX67, HEC had higher ΔG_binding_ than the STR. The most favorable ΔG_binding_ for the ligand HEC was observed with AVR-PikA, AVR-PikF, and AVR-PizT with ΔG_binding_ of -14.12, -16.12, and -13.18 kcal/mol, respectively.

**Table 2:**
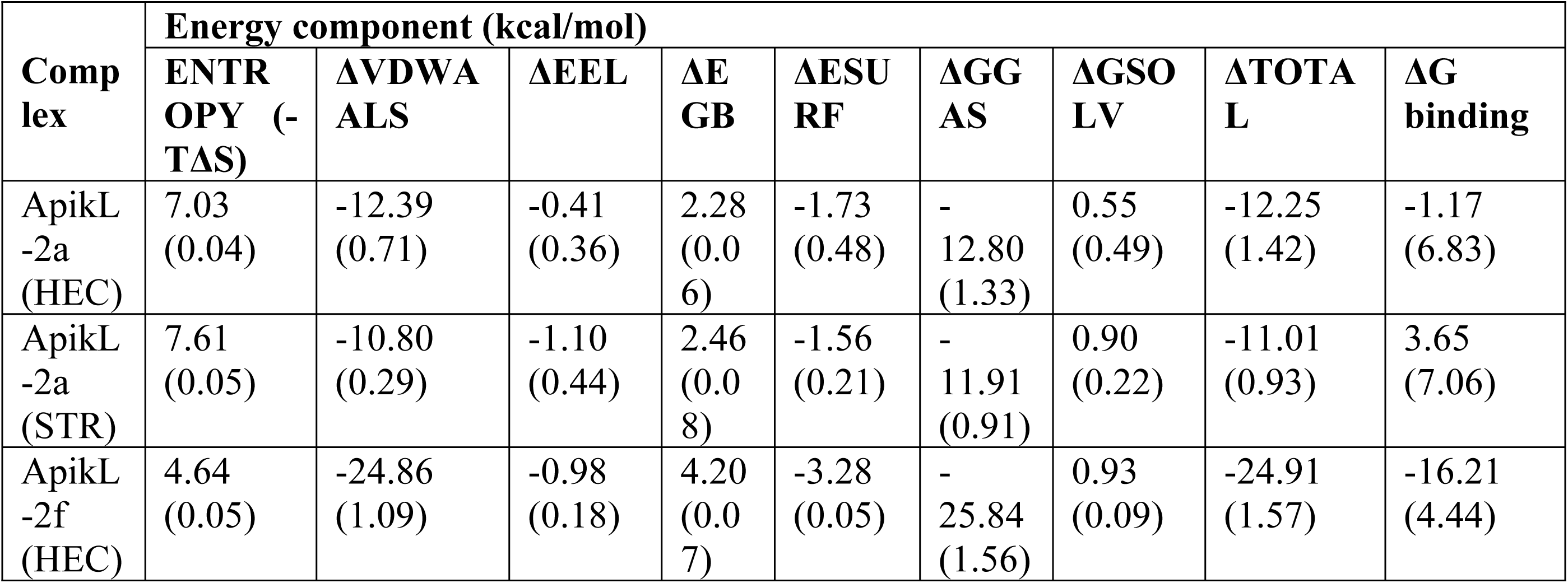
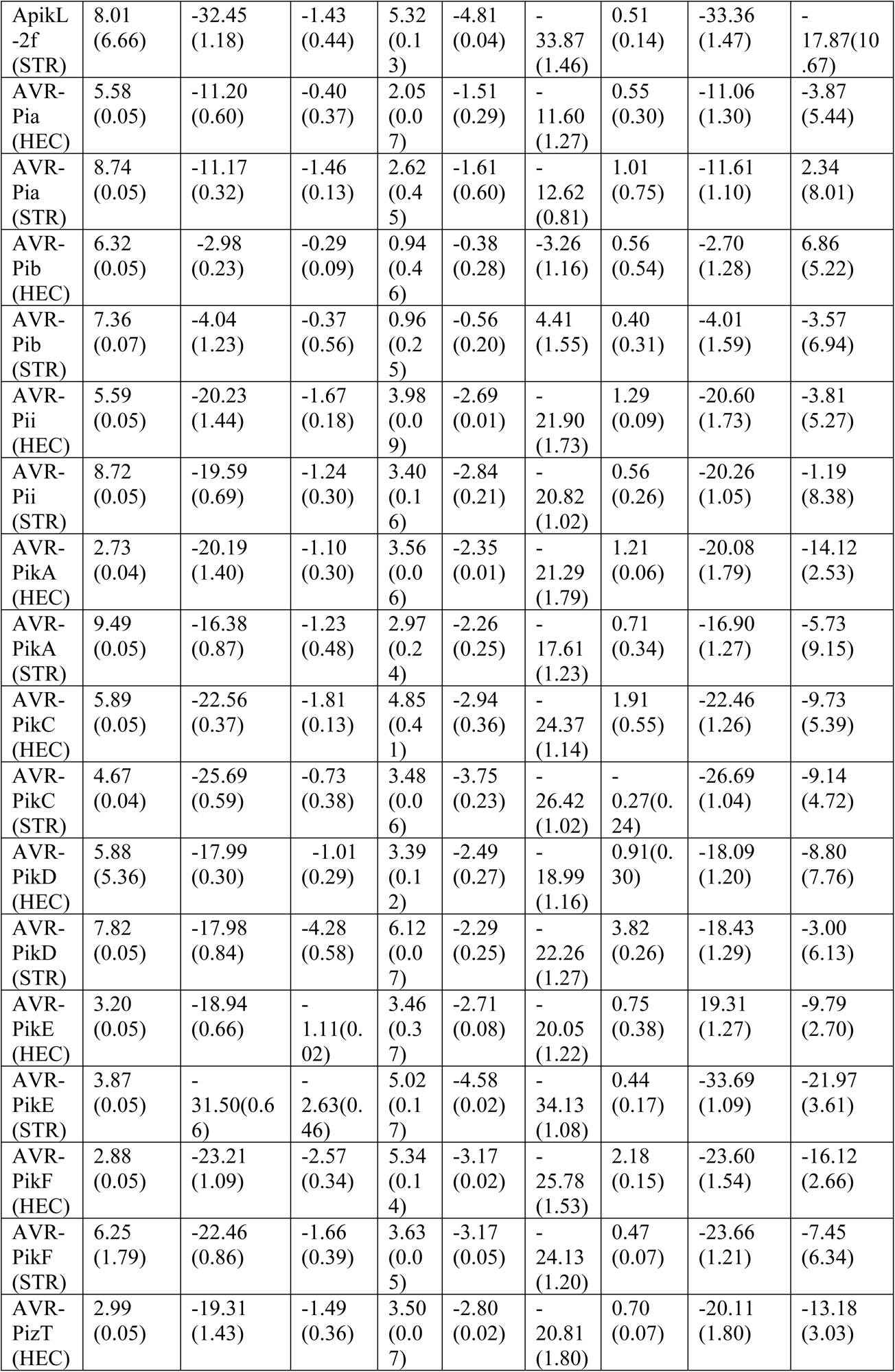
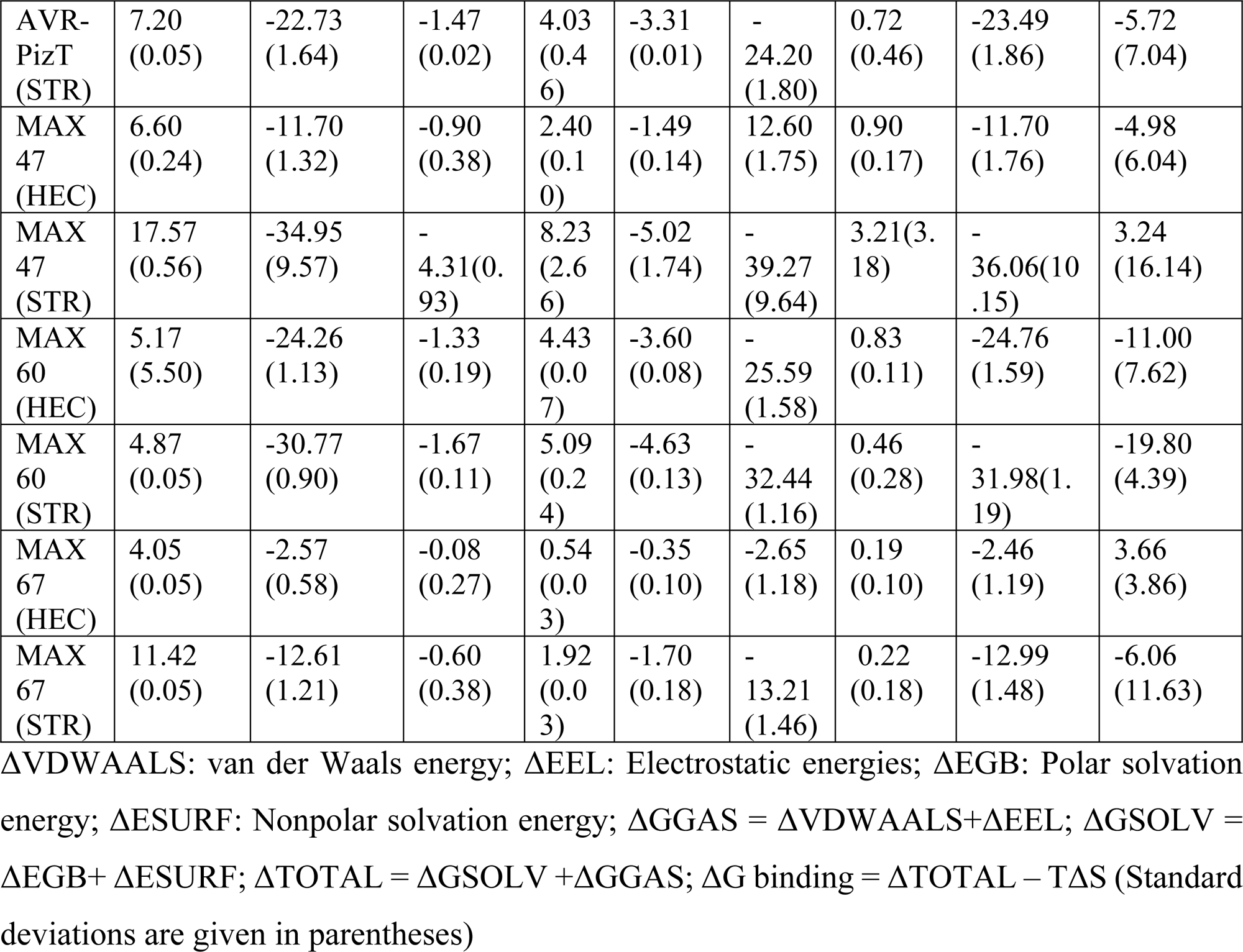
MM-GBSA results.

### 3.5. Phytochemical properties

The phytochemical properties of the compounds were analyzed using the SwissADME server. Among the 35 metabolites, a few showed negative phytochemical properties and were thus excluded from consideration as top metabolites. Hecogenin and Cucurbitacin E were further evaluated using Lipinski’s Rule of 5. **Table** 3 demonstrates that these target molecules can be considered safe for treatment. Their molecular weights range between 430 and 557 Da, and both ligands exhibit acceptable solubility. Additionally, they show good H-bond acceptor and donor characteristics, with a bioavailability score of less than 0.6.

**Table 3:** Physiochemical Properties of top metabolites.

| Properties | Hecogenin | Cucurbitacin E |
| --- | --- | --- |
| Formula | C27H42O4 | C32H44O8 |
| MW | 430.62 | 556.69 |
| Heavy atoms | 31 | 40 |
| Aromatic heavy atoms | 0 | 0 |
| Fraction Csp3 | 0.96 | 0.69 |
| Rotatable bonds | 0 | 6 |
| H-bond acceptors | 4 | 8 |
| H-bond donors | 1 | 3 |
| MR | 122.27 | 150.88 |
| TPSA | 55.76 | 138.2 |
| iLOGP | 4.06 | 3.7 |
| XLOGP3 | 4.83 | 3.25 |
| WLOGP | 4.97 | 4.19 |
| MLOGP | 4.09 | 1.68 |
| Silicos-IT Log P | 3.99 | 4.49 |
| Consensus Log P | 4.39 | 3.46 |
| ESOL Log S | -5.55 | -4.94 |
| ESOL Solubility (mg/ml) | 1.21E-03 | 6.35E-03 |
| ESOL Solubility (mol/l) | 2.80E-06 | 1.14E-05 |
| ESOL Class | Moderately soluble | Moderately soluble |
| Ali Log S | -5.73 | -5.83 |
| Ali Solubility (mg/ml) | 7.94E-04 | 8.31E-04 |
| Ali Solubility (mol/l) | 1.84E-06 | 1.49E-06 |
| Ali Class | Moderately soluble | Moderately soluble |
| Silicos-IT LogSw | -4.38 | -4.25 |
| Silicos-IT Solubility (mg/ml) | 1.78E-02 | 3.15E-02 |
| Silicos-IT Solubility (mol/l) | 4.13E-05 | 5.66E-05 |
| Silicos-IT class | Moderately soluble | Moderately soluble |
| Lipinski violation | 0 | 1 |
| Ghose violation | 1 | 3 |
| Veber violation | 0 | 0 |
| Egan violation | 0 | 1 |
| Muegge violation | 0 | 0 |
| Bioavailability Score | 0.55 | 0.55 |
| PAINS alerts | 0 | 0 |
| Brenk | 0 | 2 |
| Leadlikeness violations | 2 | 1 |
| Synthetic Accessibility | 6.7 | 6.74 |

### 3.6. Toxicity Analysis

The toxicity analysis of both top metabolites was conducted using the Deep-PK prediction server. The compounds were evaluated for potential AMES toxicity, carcinogenicity, and any irritation to the environment or human health (**Table** 4). The AMES mutagenesis probability ranged from 0.1 to 0.009, which is considered safe for the environment. The biodegradation probability ranged from 0.061 to 0.011. Additionally, the metabolites were predicted to be safe in terms of avian toxicity and carcinogenicity.

**Table 4:**
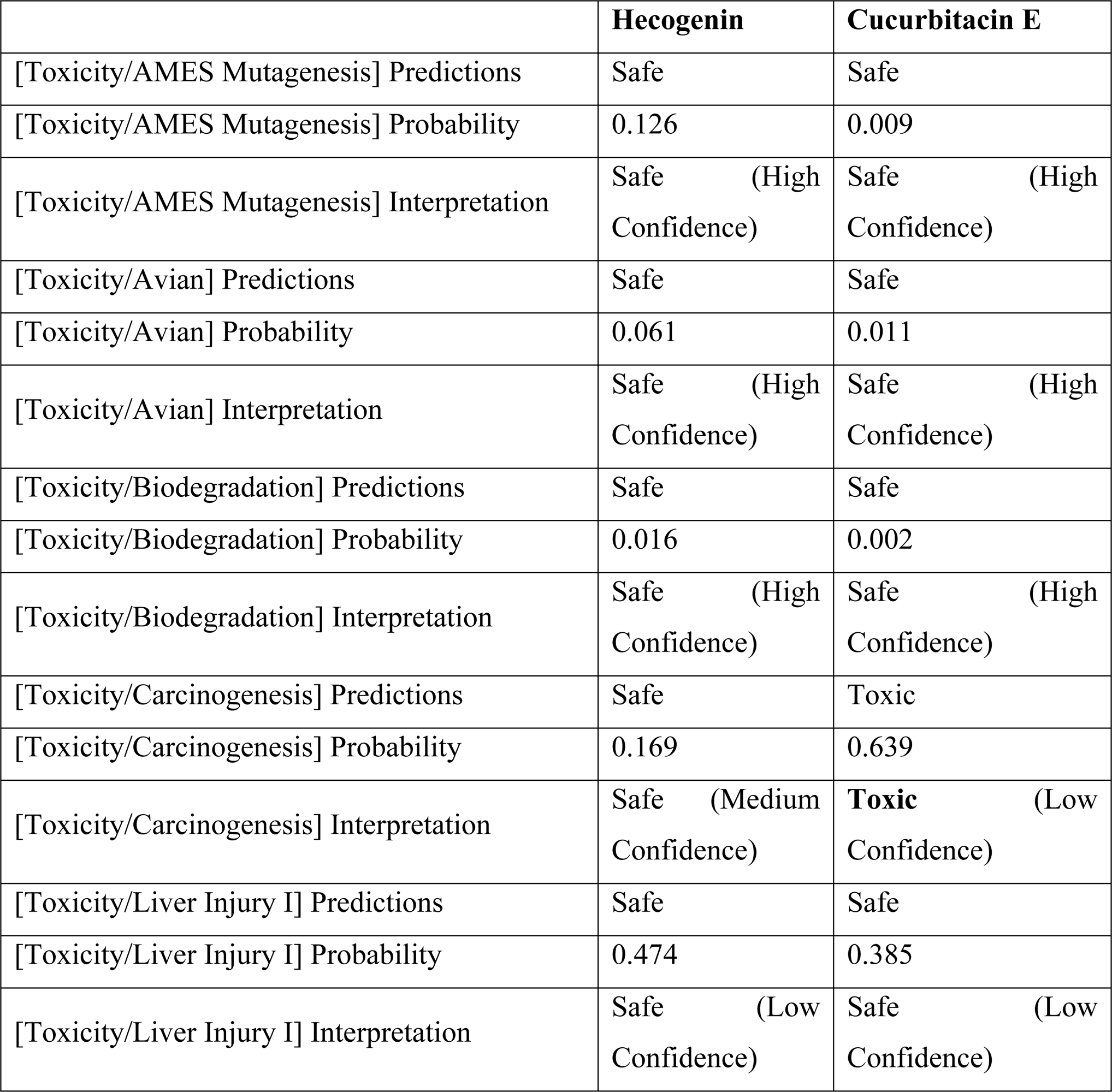
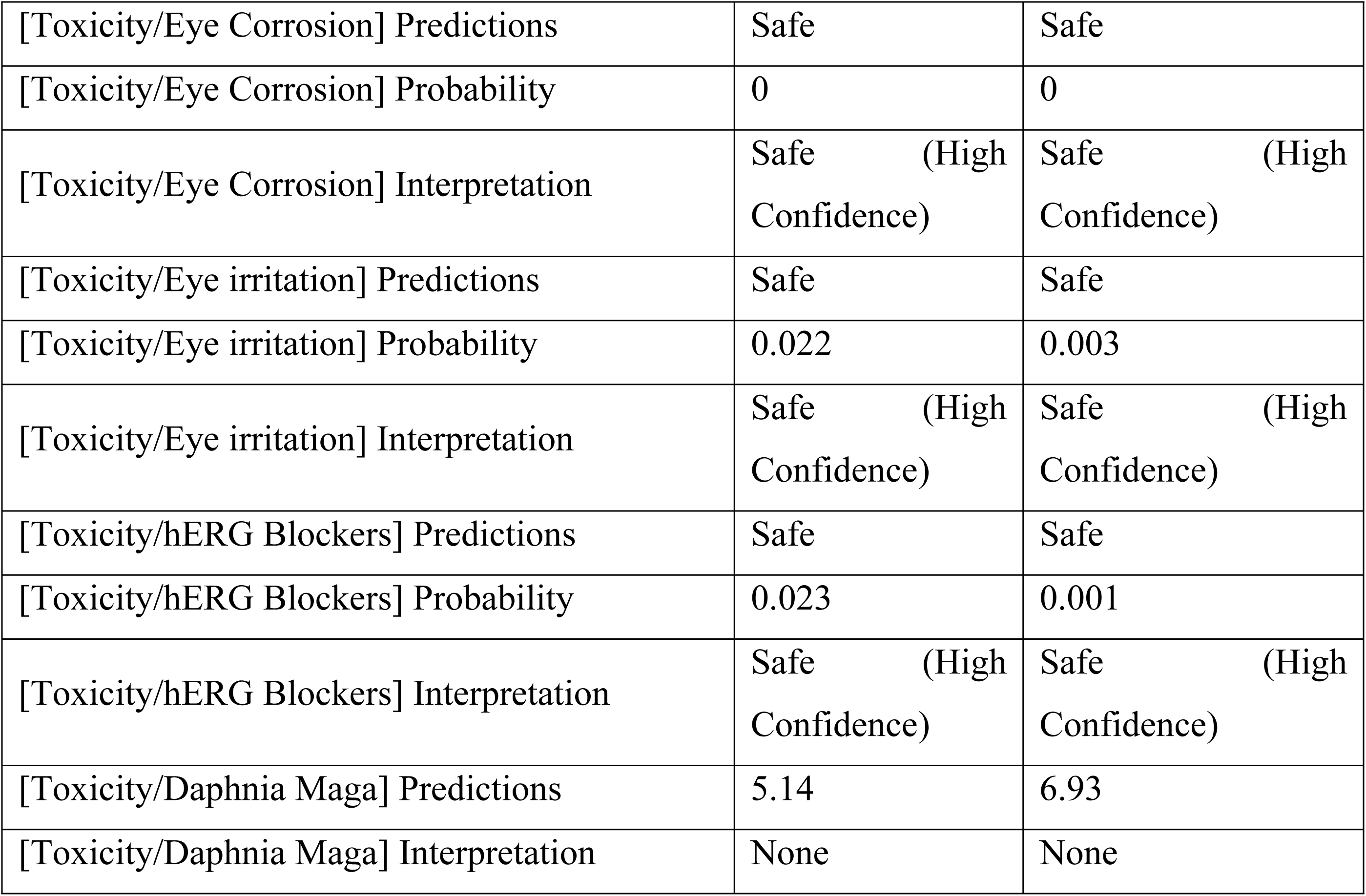
Toxicity analysis of top metabolites.

## 4. Discussion

Rice blast is a major agricultural disease causing significant financial losses worldwide. Genetic and genomic investigations have improved our understanding of the disease, with many genes implicated in its pathogenesis and development. Numerous virulence factors and signaling pathways have been identified, suggesting that the pathogen uses various techniques to infiltrate the host and subdue its defense mechanisms.

Effector proteins, including AvrPik variants and MAX proteins, have been targeted in this study because they weaken rice defenses by interacting with Pik alleles on chromosome 11. The Pik locus includes genes like Pik*, Pikm, and Pikp, which recognize AvrPik variants A to F with slight amino acid differences (Xiao et al., 2023). This interaction is mediated by the HMA domain of Pik-1, located between the CC and NB-ARC domains (Cadiou et al., 2023). Pikp-1 and Pikm-1 differ in their HMA domains, leading to distinct recognition specificities (Mukhi et al., 2020). The interaction interface has been extensively studied (De la Concepcion et al., 2018; Maidment et al., 2021). Although AvrPik-C and AvrPik-F are not recognized by any Pik alleles, AvrPik-C can interact with Pikh-HMA in vitro (Xiao et al., 2023). Advances in engineering new receptor specificities have significant implications for improving resistance against diverse pathogen strains. The AvrPib gene encodes a 75-residue protein with a signal peptide (Wang et al., 2021; Zhang et al., 2015), and M. oryzae attenuates the host ROS burst via AVR-Pii-mediated inhibition of Os-NADP-ME2 (Singh et al., 2016). MAX proteins play crucial roles in the infection process, with essential structural components like the stabilized hydrophobic core and positive-charged patch critical for AvrPib’s avirulence function (Lahfa et al., 2023; Zhang et al., 2018).

The study aimed to identify 35 compounds with potent inhibitory effects against M. oryzae for potential fungicidal applications. Among these, natural substances Hecogenin and Cucurbitacin E exhibited strong binding affinity with MAX40 and APIKL2A proteins, respectively, crucial for fungal transcription and host immunity inhibition, as revealed by molecular docking. Given M. oryzae’s reliance on these proteins for plant infection, these chemicals are promising candidates for fungicide development or as lead compounds in novel fungicide formulations targeting rice blast disease.

Fourteen essential effector proteins were refined and visualized, while thirty-five plant-derived metabolites were prepared for molecular docking analysis. Hecogenin and Cucurbitacin E consistently exhibited the highest binding affinity across all 14 effector proteins. Specifically, Hecogenin demonstrated the strongest binding energy with APIKL2A protein (-8.8 kJ/mol), while Cucurbitacin E showed the highest binding energy with MAX40 protein (-8.0 kJ/mol). Both compounds outperformed the reference metabolite, Strobilan, in binding affinity. SwissADME server analysis confirmed their favorable phytochemical properties, suggesting their potential as effective fungicides.

All the effector proteins in complexes with either HEC or STR were taken up for MD simulation studies, where the docked complexes for respective proteins were processed in MD simulations. Firstly, the objective for extensive MD simulations was to delve deeper into the stability aspects of each complex to get better insights into the binding affinities of respective ligands in each protein target. Secondly, due to the limitations of the molecular docking protocol, where the actual biological environment could not be simulated, and the protein structures were held rigid, the binding free energies might not be insightful. In this case, the MD simulations, where the actual biological environment is simulated and the overall stability of respective protein-ligand complex could be gauged, provide better insights into estimating binding affinities of ligands (Mortier et al., 2015).

The analysis of RMSD in protein backbone atoms, as well as the analysis of RMSD in ligand atoms relative to protein backbone atoms, affords insights into the overall stability of protein-ligand complexes as well as the insights into position and conformational dynamics states of ligands in the binding site during MD simulation (Magala et al., 2020). The complexes with AVR-Pikb, AVR-PikA, and AVR-PikC are stable as the RMSD in backbone atoms is well below 0.2 nm. The other complexes showing slightly higher or more significant deviations in RMSD in backbone atoms suggest the major conformational change in protein structure relative to the starting equilibrated structure. The RMSD in the ligand atoms calculated by taking the protein backbone atoms for least squares fitting provides insights into the position of ligands at the binding site and the conformational change in ligand compared to the starting structure (Kabsch, 1976). The ligands HEC and STR were quite stably bound with ApikL-2f, AVR-PikE, and AVR-PizT compared to the initial conformations of ligands. In the case of AVR-PikA, the ligand HEC seems to bind at a slightly different site than the initial binding site, as the RMSD converged after around 15 ns until the end of the simulation period.

Similarly, HEC binds stably in AVR-PikC and MAX60 at different binding sites. However, STR was found to be more stably bound at the initial binding site than the HEC. This implies that HEC might have slightly different binding interactions at an alternate binding site than the STR.

The analysis of RMSF in the protein side chain residue atoms provides further insights into structural mobility in the protein structure (Martínez, 2015). The residues in the loop region show higher RMSF due to their flexibility (Lindorff-Larsen et al., 2010), and inherently, the end terminal residues show higher RMSF. The first 50 residues in ApikL-2a and ApikL-2f are quite flexible and belong to the unstructured loop region, while other residues are quite stable to the conformational change, except terminal residues. This was observed in the complexes with both ligands. Similarly, the proteins AVR-Pia, AVR-Pib, AVR-PikA, AVR-PikC, AVR-PikD, AVR-PikE, AVR-PikF, AVR-PizT, MAX60, and MAX67 showed stable RMSF in most of the residues, except the terminal and few loop region residues. AVR-Pib in complex with STR might have unusual flexibility in the residues in the range 40-50, indicating major protein structural change. While the complexes with AVR-Pii and MAX47 having comparably high RMSF imply higher flexibility in their structures.

The radius of gyration analysis is a critical analysis that provides insights into protein compactness and stability of protein-ligand complexes (Lobanov et al., 2008). The stable and converged Rg indicates the compactness of the protein-ligand complex (Noguchi & Yoshikawa, 1998), while the deviations in Rg indicate the functional impairment of the protein-ligand complex (Blache et al., 2012). Almost all the complexes of rice-effector proteins with HEC and STR have been found compact and stable. However, significant structural changes are evident in AVR-Pii complexes, where the complex with HEC seems slightly more stable than that with STR. It is evident in the case of AVR-PikF that the complex with HEC is significantly more stable than the complex with STR which showed major functional impairments after around 45 ns simulation period. Both the complexes of MAX47 seems structurally unstable as the Rg underwent significant deviations throughout the simulation period. In the case of MAX60, the complex with HEC indicated slightly more structural changes than STR complex.

Amongst the non-bonded interactions, hydrogen bond interaction is the most influential in the binding affinity of ligands to the protein binding sites (Bissantz et al., 2010). The more the hydrogen bonds between ligands and the proteins, the better the system’s binding affinity and stability (Majewski et al., 2019). The ligands HEC and STR were seen forming a maximum of two consistent hydrogen bonds in ApikL-2a, ApikL-2f, MAX47, MAX60, and MAX67. In other complexes, the frequency of hydrogen bonds between these ligands with binding site residues was slightly lower, indicating slightly weaker binding affinities. Further, the ability of the ligands to form hydrogen bonds was observed to vary with proteins. For instance, HEC forms hydrogen bonds with Gly92 and Lys109, while STR forms with Gly97 and Leu95 in the case of ApikL-2a. Similarly, the variation in hydrogen bond formation was observed in all other complexes. HEC formed significant hydrogen bonds in AVR-Pib, AVR-PikC, and AVR-PikD complexes compared to STR.

The end-state MM-GBSA calculations, which were based on calculating the overall energies such as van der Waals, electrostatic, polar solvation, and solvent-accessible surface areas energy for protein, ligand, and protein-ligand complex, provide insights into binding affinities of ligands (Genheden & Ryde, 2015). The entropic contributions proved important in the final ΔGbinding in kcal/mol. In the case of ApikL-2a, AVR-Pia, and MAX47, the ligand HEC has a more favorable binding affinity than the STR. The favorable binding affinity of HEC might be due to lower Van Der Waal’s energy and higher electrostatic energy. Both ligands have more favorable binding affinities for HEC than STR in ApikL-2f and AVR-PikC. In the case of AVR-Pii, AVR-PiA, AVR-PikD, AVR-PikF, and AVR-PizT. However, in the complexes with AVR-Pib, AVR-PikE, MAX60, and MAX67, the ligand STR has better binding affinities than the HEC. The ligand HEC has been found to have the most favorable binding affinities compared to STR in the complexes of AVR-PikA, AVR-PikF, and AVR-PizT, where the entropic energy, van der Waals energy, and electrostatic energy contributed in the favorable binding affinities.

Afterwards, Lipinski’s rule of five was applied to examine how similar these two naturally occurring chemicals were to fungicides. The compounds that were chosen have molecular weights of more than 500 g/mol or below (430.62 g/mol and 556.69 g/mol). One of the most important factors in assessing a chemical’s fungicidal efficacy is its molecular weight.

Compared to larger molecules, molecules with a molecular weight of less than 500 are easily transported, dispersed, and absorbed by the cell membrane (Leeson, 2012). A compound’s ability to flow through biomembranes is also indicated by positive LogP values, where a value of less than 5 is considered acceptable (Chang et al., 2004; Refsgaard et al., 2005).

It was found in the study that lipophilic chemicals can easily pass through the cell membrane and bond with biomolecules, blocking essential metabolic enzymes. The LogP values of the chosen natural compounds were within the range, making them suitable for passing through the fungus’s cell membrane. It was observed in this investigation that one of the compounds had a molecular weight that was higher than the predicted limit stated in Lipinski’s rule of five. This little increase in molecular weight may not have a major impact on the natural compound’s transportation and diffusion. Several FDA-approved medications have been demonstrated to have molecular masses more than 500 g/mol (Mullard, 2018). Additionally, there were fewer than five hydrogen bond donors and fewer than ten hydrogen bond acceptors.

For toxicity analysis, the Deep-PK prediction server was used. It was revealed that potential fungicide candidates are considered safe. The mutagenesis probability for both compounds ranged between 0.009 and 0.126, and avian toxicity predictions were within 0.011 to 0.061, indicating that their use will not cause any harmful effects. However, cucurbitacin E is relatively more toxic than hecogenin, and this carcinogenesis probability could be addressed through enhanced modification of the drug during preparation. The main concern for drug design is the hERG blocker, which is also considered safe during toxicity assessment.

## 5. Conclusion

This study identified Hecogenin and Cucurbitacin E as potent antifungal agents against *M. oryzae* through extensive bioinformatics analysis, including molecular docking and dynamics simulations. These compounds demonstrated strong and consistent binding affinities with 14 effector proteins, outperforming the reference compound, Strobilurin. These findings suggest that Hecogenin and Cucurbitacin E hold promise as therapeutic agents for combating rice blast disease. Future in vitro and in vivo studies are necessary to validate these results and explore the practical applications of these metabolites. Further optimization may enhance their efficacy and safety for potential use in agricultural practices.

